# Landscape and dynamics of the transcriptional regulatory network during natural killer cell differentiation

**DOI:** 10.1101/572768

**Authors:** Kun Li, Yang Wu, Young Li, Qiaoni Yu, Zhigang Tian, Haiming Wei, Kun Qu

**Author notes:** Correspondence: Kun Qu and Haiming Wei. Co-first authors (Kun Li, Yang Wu). **Contact Information**: Kun Qu Ph.D., Division of Molecular Medicine, Hefei National Laboratory for Physical Sciences at Microscale, the CAS Key Laboratory of Innate Immunity and Chronic Disease, CAS Center for Excellence in Molecular Cell Science, School of Life Sciences, University of Science and Technology of China. Hefei, Anhui, China, 230027.

## Abstract

Natural killer (NK) cells are essential in controlling cancer and infection. However, little is known about the dynamics of the transcriptional regulatory machinery during NK cell differentiation. In this study, we applied assay of transposase accessible chromatin with sequencing (ATAC-seq) technique in a self-developed *in vitro* NK cell differentiation system. Analysis of ATAC-seq data illustrated two distinct transcription factor (TF) clusters that dynamically regulate NK cell differentiation. Moreover, two TFs from the second cluster, FOSL2 and EGR2, were identified as novel essential TFs that control NK cell maturation and function. Knocking down either of these two TFs significantly impacted NK cell transformation. Finally, we constructed a genome-wide transcriptional regulatory network that provides an understanding of the regulatory dynamics during NK cell differentiation.

## Introduction

Natural killer (NK) cells are innate lymphocytes, and as the name suggests, NK cells secreting inflammatory cytokines and directly killing infected cells represent the early defense line during pathogen invasion, which surveys the environment and to protect the host from infection or cancer cells. [1]. Additionally, NK cell-based immunotherapy has become an emerging force in cancer treatment and will play an essential role in the future treatment [2–6]. For instance, adoptive NK cell immunotherapy has become increasingly popular because it induces graft-versus-leukemia effects without causing graft-versus-host disease in patients [7]. Therefore, major efforts are currently underway to fully utilize the anti-tumor properties of NK cells in the clinic. In addition, a variety of methods include the development of NK cell expansion protocols to establish a microenvironment that favors NK cell activity, redirect the activity of NK cells on tumor cells and release the inhibitory signals that limit NK cell function [8]. On the other hand, many researchers used umbilical cord blood (UCB) CD34^+^ cells to produce abundant NK cells [9–18] that were used for clinical application without feeding cells by adding various cytokines [10]. In our previous work, we have developed a method to obtain sufficient functional NK cells by simply adding a mixture of cytokines and providing a mechanism by which NK cells can be used to treat leukemia [19]. The use of NK cells for immunotherapy relies on a great number of NK cells with optimal cytotoxic activity [4]; therefore, a comprehensive understanding of the regulatory circuits during NK cell differentiation is particularly important for boosting the efficacy of clinical treatments. However, the mechanisms underlying the differentiation of the NK cells are not well understood. Current studies have shown that transcriptional factors (TFs) play an essential role in driving NK cell maturation, and many TFs have been well studied in this process [1]. Additionally, it is known that different TFs play various roles at distinct stages of differentiation [1]. For example, *PU.1* is a TF that is known to drive hematopoietic stem cell (HSC) differentiation into the earliest myeloid and lymphoid progenitors [20], whereas *T-bet* is an essential TF in the control of NK cell maturation and IFNγ secretion regulation [21]. However, how TFs work in concert to enforce the NK cell phenotype is not clear. In our work, we investigated the landscape and dynamics of the transcriptional regulatory network during NK cells differentiation from cord blood CD34^+^ HSCs based on a newly developed epigenomic profiling technique named ATAC-seq [22].

ATAC-seq has been widely used to profile the epigenetic landscapes of cells at specific stages of interest and thereby delineate the underlying regulatory mechanisms of gene expression. For example, a previous report used ATAC-seq to help identify an exhaustion-specific enhancer that regulates PD-1 expression and thereby elucidated the regulatory mechanism of gene expression in exhausted CD8^+^ T cells [23, 24]. Alternately, ATAC-seq analysis of pure cancer cell populations of human small cell lung cancer (SCLC) identified a novel TF *Nfib* which is necessary and sufficient to drive the metastatic ability of SCLC cells [25]. ATAC-seq has also enabled researchers to track the epigenomic states changes in patient-derived immune cells [26], and to survey how the personal regulomes of the cutaneous T cell lymphoma patients determines their responses towards histone deacetylase inhibitors (HDACi) anti-cancer drugs [27]. More recently, ATAC-seq was applied to better understand the controlling of NK cells in innate immune memory during infection [28], illustrating the importance of the topic as well as the power of the technique used in this study. Here, we have developed a systematic method to characterize the chromatin accessibility and construct regulatory network dynamics during NK cell differentiation based on ATAC-seq. Motif and enrichment analysis from HOMER [29] and Genomica [26] show that many TFs play important roles in NK cell differentiation, and by integrating gene expression profiles from our previous study [19], we identified two novel transcription factors, *FOSL2* and *EGR2*, that are essential for driving NK cell maturation.

## Results

### Landscape of DNA accessibility during NK cell differentiation

To elucidate the regulatory networks during NK cell differentiation, we developed a culture procedure to obtain differentiated NK cells from UCB CD34^+^ cells using a cocktail of cytokines [19]. The differentiation process requires 35 days. Interestingly, after culturing of cord blood stem cells for 3 weeks, the proportion of NK cells rapidly increased from 5% on day21 to approximately 60% on day 28, and peaking at close to 100% on day 35 (**Figure 1A** and **Figure S1A**) [19]. To elucidate the molecular mechanism and regulatory network underlying NK cell differentiation, we interrogated the landscape of chromatin accessibility using ATAC-seq at 8 different time points, with 2 replicates each along the process (**Figure 1B**). Multiple bioinformatics analysis methods (see **Materials and Methods**) were then applied to obtain the differential accessible chromatin sites, enriched transcription factors (TFs) and genome-wide regulatory elements. At least 50,000 cells were obtained from each sample, and on average, resulted more than 78 million reads with a total of 1260 million measurements (**Table S1**). From this dataset, 143,570 DNA accessible sites were identified (P<10^−7^, FDR<10^−7^). Quantitative analysis indicated that the dataset was of high quality with a strong signal to background noise ratio and expected fragment length distribution (**Figure S1B-E**). Moreover, the dynamics of the chromatin accessibilities around the known surface marker genes were consistent with changes of their gene expressions during NK cell differentiation (**Figure S1D**) [30, 31]. Pearson correlation analysis on the biological replicates at each time point also showed that our ATAC-seq profiles were highly reproducible **(Figure S2A-B)**. Within all the accessible sites, only a very small portion (6.48%) were conserved across all stages **(Figure S3A)**, while the majority of peaks changed over time, suggesting significant chromatin dynamics during the NK cell differentiation. Peaks that were detected at all stages were enriched in gene ontology (GO) terms such as RNA transcription and other functions to maintain basic physiological activities **(Figure S3B)**. In contrast, stage-specific peaks, especially peaks that emerged at later stage of the process, were highly enriched in immune relevant GO terms, suggesting the activation of critical genes that govern NK cell function.

**Figure 1.**
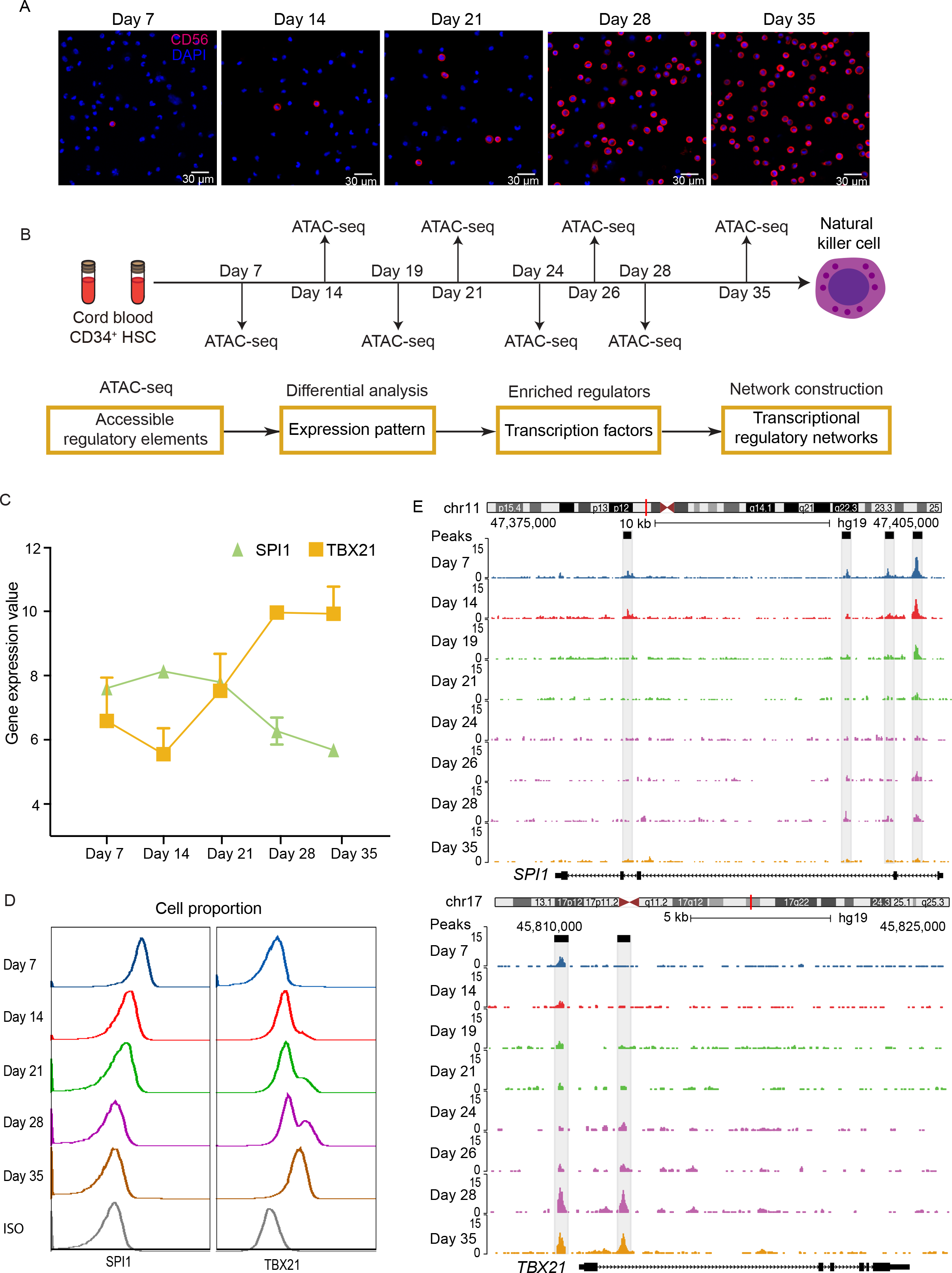
Landscape of DNA accessibility during NK cell differentiation. **A** Confocal microscopy images of membrane CD56 (red) in the cultured cells at days 7, 14, 21, 28 and 35. Scale bars, 30 µm. Nuclei are stained with DAPI. **B** Schematic representation of the overall experimental design of this study. Gene expression and chromosome opening at different time points were assessed using microarray and ATAC-seq data respectively. The bioinformatics pipeline for data analysis is shown in the bottom. **C** The gene expression profiles of *SPI1* and *TBX21* at different stages of NK cell differentiation. **D** Flow cytometric measurement of PU.1 (*SPI1*-encoded protein) and T-bet (*TBX21*-encoded protein) expression using in cultured cells during a 35-day time course. **E** Normalized ATAC-seq profiles of the *SPI1* (top) and *TBX21* (bottom) gene loci at different stages during NK cell differentiation. These two genes are known to regulate NK cell differentiation, and ATAC-seq signals were obtained from the UCSC genome browser.

For instance, several TFs are known to regulate NK cell differentiation, such as SPI1 and TBX21. The former is a NK cell repressor and the latter an activator. Gene expression profiles from our previous study [19] indicated down-regulation of the SPI1 gene and up-regulation of the TBX21 gene during NK cell differentiation (**Figure 1C**). The protein levels of these two genes obtained from flow cytometric experiments suggested the same trends (**Figure 1D**). ATAC-seq revealed two regulatory elements at the promoters and intronic enhancers around each of these two genes (**Figure 1E**). Chromatin-accessible sites around the *SPI1* gene clearly became unavailable and those around the *TBX21* gene gradually became available during the process, supporting the notion that the epigenetic dynamics of key regulators are consistent with their corresponding gene and protein expressions. The consistency between the epigenetic and gene expression profiles of transcription factors, such as GATA3 and EOMES, which are known to drive NK cell differentiation, are shown in (**Figure S1E**), indicating the feasibility of predicting transcriptional regulatory networks from ATAC-seq profiles.

### Epigenomic signatures of different stages during NK cell differentiation

To determine the differences in regulatory DNA activity among different stages during NK cell differentiation, we applied pairwise comparisons of the ATAC-seq signals between the corresponding samples, together with intrinsic analysis [32], a method that highlighted elements that varied in accessibility across individuals but not between repeat samples from the same individual. We discovered 6401 peaks of differential DNA accessible sites across the genome, and identified three distinct clusters of chromatin accessible sites via unsupervised hierarchical clustering (**Figure 2A, Figure S4A, B**). Principal component analysis (PCA) of all the samples also illustrated three distinct clusters, which were consistent with the time process of NK cell differentiation representing the early, intermediate and late stages of the entire process (**Figure S4C**). Gene ontology analysis of these peaks was performed in GREAT [33]. Cluster I comprises of 1584 elements that were more accessible at the early stage (days7∼21) of differentiation. Peaks in cluster I were enriched in the GO terms of metabolic processes, with a certain level of significance (**Figure 2B**,10^−8^<P<10^−4^), keeping cells alive, proliferation and prepare to differentiate. Cluster II comprises 4817 elements, which were highly accessible at the late stage (days 24∼35) of NK cell differentiation. GREAT analysis revealed that peaks in cluster II were strongly enriched (P<10^−50^) for immune-relevant GO terms, such as immune system process, immune response and immune system development among others (**Figure 2B**), indicating that the epigenetic states of functional immune cell-specific genes were activated throughout the process and then drove progenitor cells differentiation toward NK cells. Cluster III consisted of only 356 peaks, which showed no enrichment for GO terms and were therefore neglected in downstream analysis.

**Figure 2.**
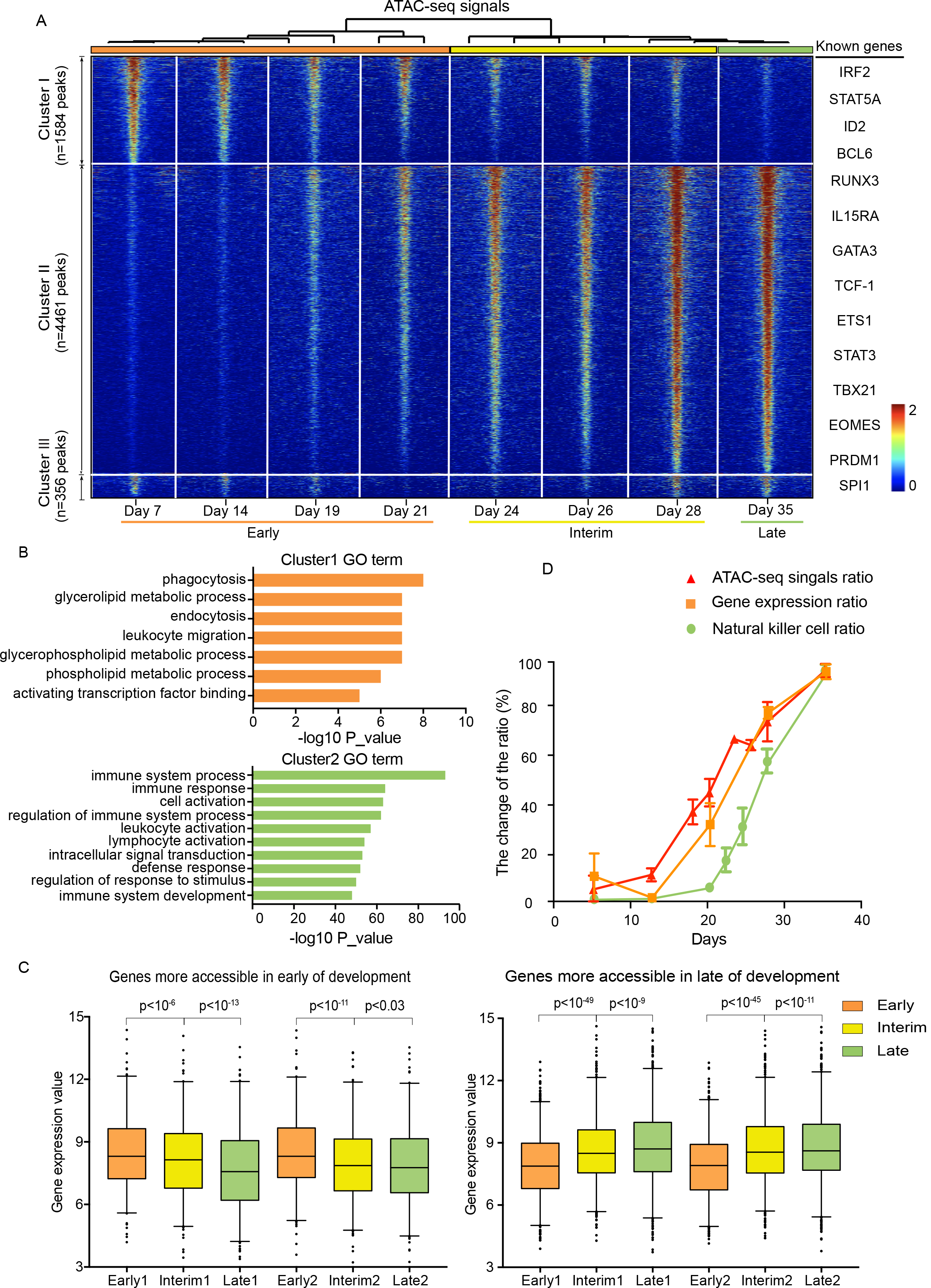
Epigenomic signatures of different stages during NK cell differentiation. **A** Heatmap of 6401 differentially expressed regulatory elements during NK cell differentiation. Each column is a sample, and each row is a peak. Samples and peaks are organized by two-dimensional unsupervised hierarchical clustering. Color scale showing the relative ATAC-seq peak intensity centered by each peak summit. Top: samples at all time points were categorized into three groups, early stage: days 7∼21 (orange); interim stage: days 24∼28 (yellow) and late stage: days 28∼35 (green). Samples from the same cluster are labeled with the same color. Left: differential peaks are categorized into 3 clusters. **B** The top ten most significant gene ontology (GO) terms enriched in cluster I and cluster II peaks. **C** Box-plots of the mRNA expression levels of the genes that are more accessible in early, interim and late stages during NK cell differentiation. 1,2 represents biological replicates 1 and 2, respectively. P-values were estimated from Student’s t-test. **D** The changes in ATAC-seq signals (red), gene expression (orange) and number of NK cells (green) at different time points during NK cell differentiation.

We next examined whether the DNA accessibility chromatin signatures at different stages correlated with those of the corresponding gene expressions. By comparing the ATAC-seq profiles with the genome-wide microarray data during NK cell differentiation, we found that gene loci that gained chromatin accessibility were significantly increased in the gene expression level (P<10^−6^), while gene loci that lost chromatin accessibility had decreased expression (P <10^−9^), indicating a high correlation between epigenetic and RNA profiles (**Figure 2C**).

Chromatin structure and epigenetic modifications regulate gene expression [34], however, the chronological order of the dynamics of the chromatin accessibility, gene expression, and cell phenotype have not yet been well studied. Here, we integrated the ATAC-seq signals, microarray profile and percentage of NK cell counts to examine the temporal dynamics of these three features. Interestingly, we noticed that the accessibility of mature NK cell-specific peaks (cluster II in **Figure 2A**) started to open up on day 14; and the expression of NK cell specific genes was turned on approximately two days later, while the percentage of NK cell started to grow after day 21 (**Figure 2D**). These results suggested a clear chronological order of the changes in chromatin structure, gene expression and cell phenotype during NK cell differentiation.

### Transcription factor occupancy network during NK cell differentiation

Transcription factors bind to cognate DNA sequences with patterns termed motifs, and therefore, we can predict TFs occupancy on chromatin by DNA accessibility data from ATAC-seq, and use it to create regulatory networks [26]. To identify potential drivers of NK cell differentiation, we applied HOMER and used both the default (**Figure 3A**) and setting background (**Figure S5A**) search modes (see **Materials and Methods**) to search for TFs that were enriched at accessible elements in cluster I and cluster II suggesting that NK cell differentiation and maturation require a variety of transcription factors. TFs enriched at cluster I peaks are potential regulators at the early stage of NK differentiation, while those enriched at cluster II peaks could be critical regulators at the late stage. We found several TF families that were significantly enriched (P<10^−10^), and many of which are important known TFs of NK cells differentiation, such as the RUNX and ETS1/PU.1 families [20, 35] and CEBP [36], supporting the reliability of this method to detect critical regulators. For instance, PU.1 is widely expressed and controls multiple stages of bone marrow and lymphocyte differentiation in a variety of hematopoietic-derived lineages [37–41], and a reduction in the number of NKPs and iNKs has been detected in chimeric mice [20], indicating that PU.1 may play an important role during the early stages of NK cell differentiation. Several known TFs are also enriched at the late stage, such as RUNX [35], E2A [42], T-bet [21], STAT5 [43], and EOMES [21]. The most enriched motif in cluster II was the RUNX family: RUNX1, RUNX and RUNX2 (**Figure S5A**), which are key regulators of lymphocyte lineage-specific gene expression [44].

**Figure 3.**
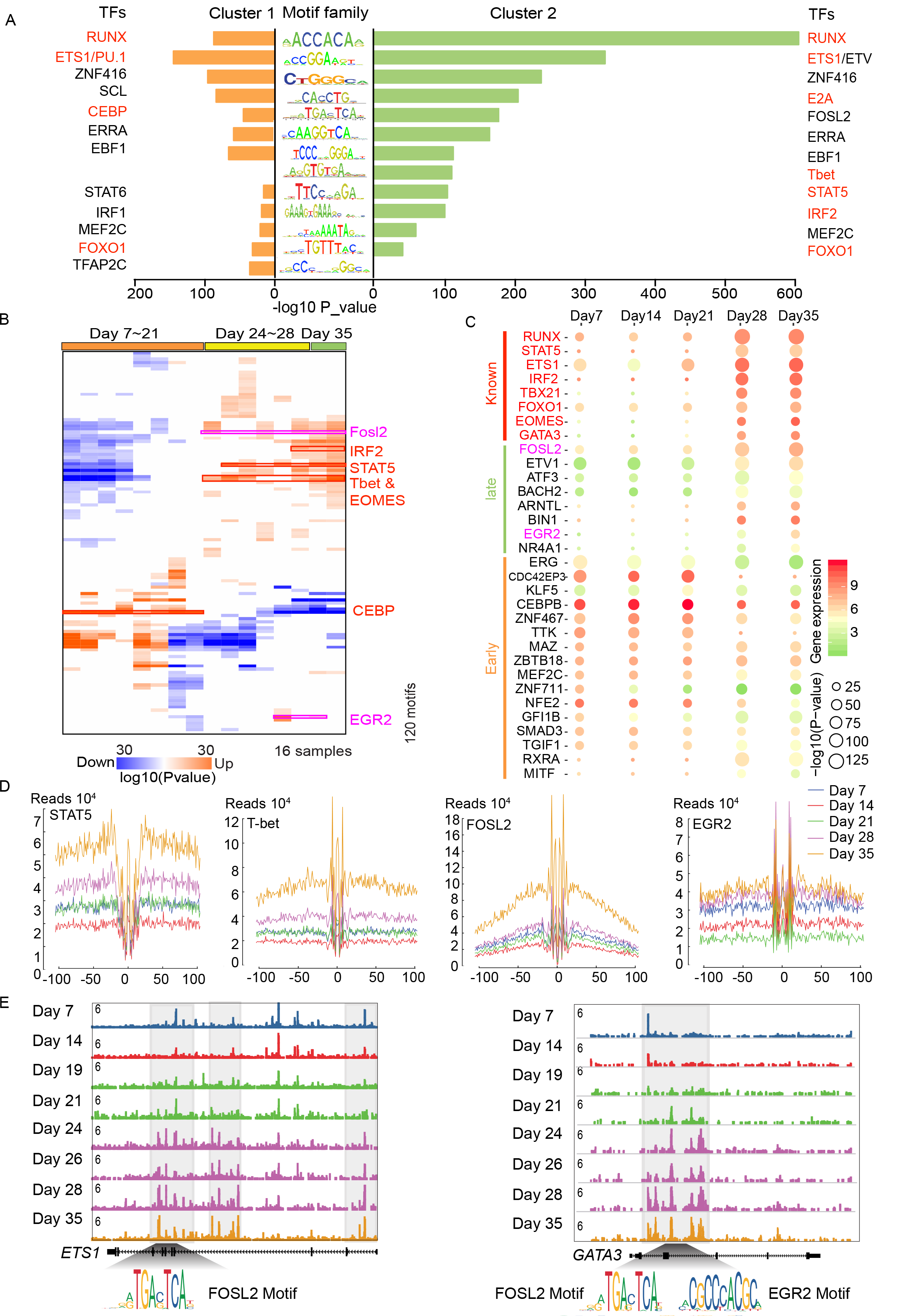
Transcription factor occupancy network during NK cell differentiation. **A** TF motifs enriched in cluster I (left) and cluster II (right) peaks, with enrichment P-values estimated from HOMER. TFs known to regulate NK cell differentiation are colored red. **B** Enrichment of known TF motifs in differential accessible elements in all samples. Each row is a motif, and each column is a sample. Values in the matrix represent the significance level in terms of the -log(P-value) of the enrichment estimated from Genomica. Red indicates that the motif is enriched in the corresponding sample, and blue indicates depletion. Red box indicates known key TFs that regulates NK cell development, the pink box indicates the new TF whose function will be experimentally tested later in Figure 4. **C** Prediction of TFs that may regulate NK cell differentiation. TFs known to regulate NK cell differentiation are shown at the top, TFs predicted to regulate early stages of the differentiation process are shown in the bottom, and those that regulate the late stage are shown at the middle. The color of each circle represents the expression level of the gene encoding the corresponding TF, while the size of the circle represents the significance of the motifs estimated by P-values. **D** Visualization of the ATAC-seq footprint for STAT5, TBET, FOSL2 and EGR2 motifs during five stages of NK cell differentiation: day 7 (blue), day 14 (green), day 21 (red), day 28 (cyan), day 35 (purple). ATAC-seq signals across all these motif binding sites in the genome were aligned on the motif and averaged. **E** Normalized ATAC-seq profiles of the *ETS1* (left) and *GATA3* (right) gene loci at different stages during NK cell differentiation. These two genes are known to regulate NK cell differentiation, and ATAC-seq signals were obtained from the UCSC genome browser. Bottom: The motif of FOSL2 and EGR2.

Motif searches of differential ATAC-seq peaks could potentially provide information about the transcriptional regulatory network, but one caveat of this method is unable to distinguish between similar motifs of binding TF family members. Thus, we sought to apply the Genomica’s module map algorithm and “TF footprint” analysis to better predict TF occupancy on accessible sites by integrating the ATAC-seq and gene expression microarray profiles (**Figure 3**).

From the Jasper database we obtained 242 vertebrate TF motifs [45]. We then used Genomica [27] producing a time points-specific TF occupancy network (**Figure 3B**). This analysis revealed distinct patterns of TF access to DNA at different time points. Many known TFs were enriched, for instance, STAT5, which is an IL-15 downstream signaling molecule and is indispensable throughout the lifetime of NK cells. Deficiency of STAT5a/b has been reported to result in complete elimination of NK cells, which demonstrates the important and non-redundant effects of STAT5 [43]. The JAK-STAT pathway has also been shown to be an important signaling pathway used by various cytokines and growth factors [46]. The interferon regulatory factor (IRF) family regulates a variety of processes, including hematopoietic development, tumorigenesis, host immunization and pathogen defense [47, 48]. IRF2 is required to maintain the normal differentiation of NK cells in a cell-intrinsic manner [49, 50]. And IRF2-deficient NK cells showed reduced levels of mature markers and IFN-γ production during stimulation [49, 50]. T-bet and EOMES are members of the T-box family and are known to control different aspects of NK cell differentiation and maturation. [21, 51, 52].

In addition, several novel TFs were also enriched, namely, FOSL2 and EGR2, and are potential regulators of NK cell differentiation. These TFs were assessed along with the other enriched TF families from the motif analysis to identify which gene in the family was expressed or differentially expressed during NK differentiation to further filter out true regulators. At each time point, we plotted both the expression value (colored from red to green to represent high to low expression in the gene microarray profile) and the motif enrichment score (shown by the circle size representing the –log(P-value) of the enrichment) in the same figure (**Figure 3C**), and we observed that the known regulators were highly expressed and enriched at different stages, same as the genes *FOSL2* and *EGR2*. By integrating results from the above three analyze (**Figure 3A-C**), we predict FOSL2 and EGR2 as potential regulators.

DNA sequences that are directly occupied by DNA-binding proteins are protected from transposition, and therefore the resulting sequence “footprint” reveals the presence of a DNA-binding protein at each site, analogous to DNase digestion footprints. TF “footprint” analysis of our ATAC-seq profile provided further evidence of direct occupancy of a TF candidate on genomic DNA and thus refined the prediction of potential regulators. We illustrated the “footprints” of 2 known regulators, STAT5 and T-bet, and observed higher DNA accessibility and deeper “footprints” flanking their motifs in the late compared with the early stage during NK cell differentiation (**Figure 3D**). Similarly, “footprints” of the TFs FOSL2 and EGR2 were also deeper and more accessible at the late stage, suggesting not only that the motifs of these 2 TFs were enriched at stage-specific peaks but that they were most likely physically bound to those accessible chromatin sites, indicating that they are functional regulators of NK differentiation (**Figure 3D**). Overall, the results from footprint analysis were also consistent with that from the HOMER and Genomica’s motif enrichment analysis.

By combining the TF motif and “footprint” analysis, we can theoretically predict genes that are regulated by any TF of interest. We performed this analysis on the TFs RUNX, *T-bet*, FOSL2 and EGR2 and integrated the gene expression profiles at each time point (**Figure S5B**, **Table S2**). We found that genes regulated by each of these TFs were also highly expressed at late stages and were significantly enriched in GO terms of immune system construction and other related functions. Transcription factors ETS1 was reported to drive early stages of NK cell differentiation [53], and we found that there was a FOSL2 binding site in a dynamical accessible sites on the gene body of *ETS1*, suggesting FOSL2 might regulate NK cell differentiation through *ETS1* (**Figure 3E**). Transcription factors GATA3 was found regulating liver-specific NK cells, IFN-γ production, and T-bet expression in mice [54]. Similarly, we found that both FOSL2 and EGR2 bind to the gene body of *GATA3* (**Figure 3E**), suggesting that the TFs FOSL2 and EGR2 may also regulate NK cell differentiation through regulation of *GATA3*.

### FOSL2 and EGR2 are necessary for NK cell differentiation

*Fosl2* belongs to the *Fos* gene family, which encodes leucine zipper proteins that combines to the TF complex AP-1 in the form of a dimer with JUN family. Thus, FOS proteins have been suggested as key regulators of transformation, differentiation and cell proliferation. *FOSL2* (FOS-like 2) is a protein-coding gene. The GO annotations of *FOSL2* include sequence-specific DNA binding, transcription factor activity and RNA polymerase II specific DNA binding. A previous report has shown that *FOSL2* is constitutively expressed in adult T-cell leukemia (ATL) and up-regulates CCR4 and promotes ATL cell growth, together with JUND [55]. *EGR2* (early growth response 2) is also a protein-coding gene that contains three tandem C2H2-type zinc fingers. GO annotations of this gene include ligase activity, sequence-specific DNA binding and transcription factor activity. Previous reports have shown that *Egr2* regulate T-cell and B-cell function in homeostasis and adaptive immune responses by controlling inflammation and promoting antigen receptor signaling [56, 57].

Since their regulatory functions in NK cell differentiation have not been well characterized, we performed loss-of-function experiments to assess the effects of *FOSL2* and *EGR2* on NK cell differentiation. We first validated their expression levels with real-time PCR, and we found that their expression gradually increased in later stages of NK cell differentiation, consistent with our bioinformatics exploration (**Figure S6A**). Subsequently, we infected the cultured cells with TF-specific shRNA- and control shRNA-containing lentivirus, respectively, which are represented by GFP expression (**Figure 4A**). We then sorted the GFP-positive cells and observed dramatically reduced FOSL2 and EGR2 expression levels (**Figure 4B**), suggesting positive targeting of the TF-specific shRNA. During NK cell differentiation, we observed a nearly 30% reduction in the proportion of differentiated NK cells in GFP-positive cells at days 28 and 35 (**Figure 4C-D**). However, in the non-transfected (GFP-negative) cells, the proportion of differentiated NK cells was not affected at the same time (**Figure 4E**-F). These results indicate that knockdown of *FOSL2* and *EGR2* expression, but not viral infection, inhibit NK cell differentiation, suggesting that *FOSL2* and *EGR2* are necessary for NK cell differentiation. We then tested another important marker of NK cell maturation CD11b [1], and found that it was significantly reduced in GFP-positive NK cells, suggesting that *FOSL2* and *EGR2* might affect NK cell maturation (**Figure 4G**). Overall, we predicted these two TFs FOSL2 and EGR2 as key regulators based on ATAC-seq and gene microarray profile analysis, and then experimentally verified that they were indeed necessary for normal NK cell differentiation.

**Figure 4.**
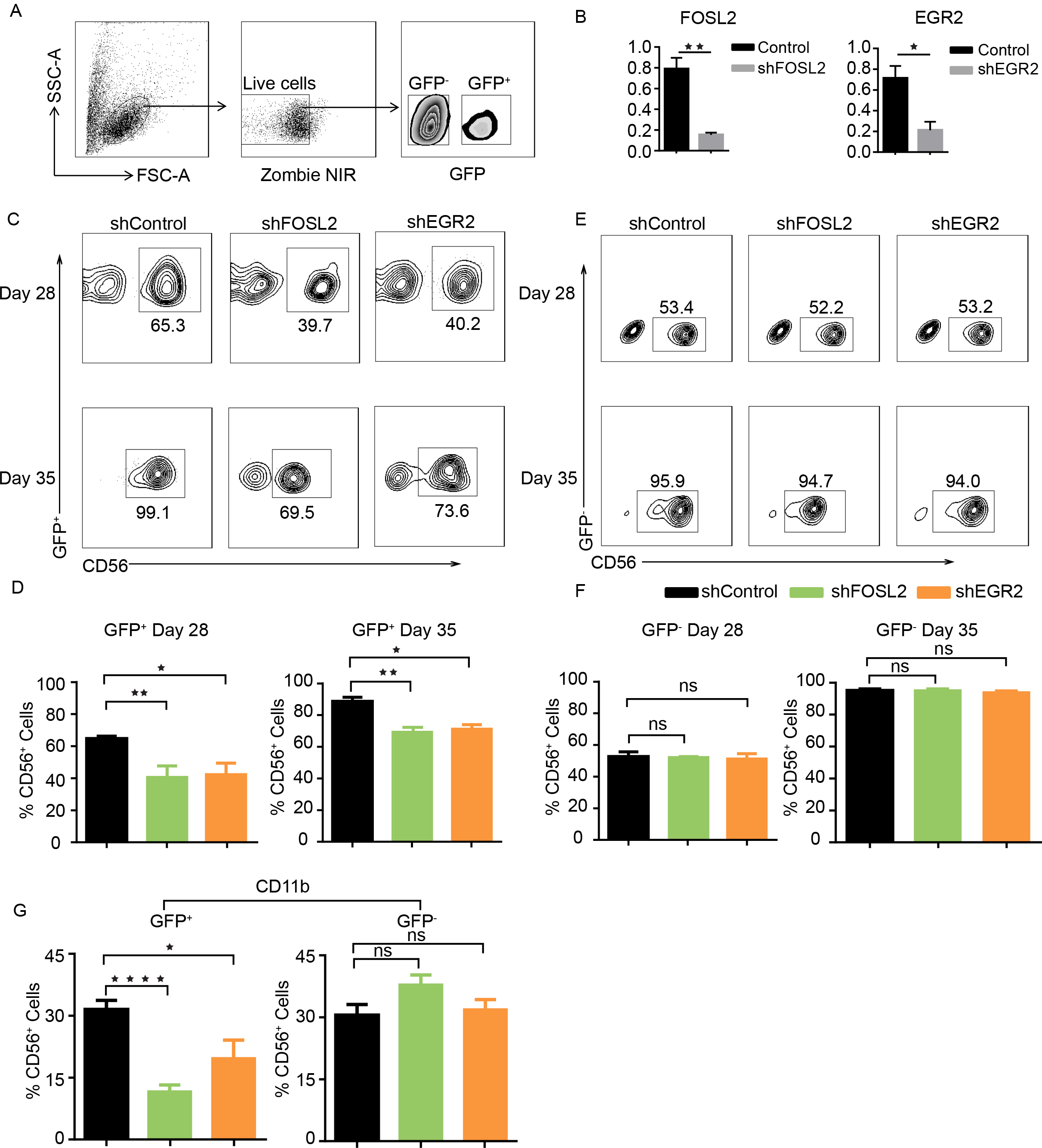
FOSL2 and EGR2 are necessary for NK cell differentiation. **A** The gating strategy for cultured cells transduced with lentiviruses expressing either shFOSL2, shEGR2 or control shRNA via detection of GFP expression. **B** Knockdown efficiency of *FOSL2* and *EGR2* with shFOSL2 and shEGR2. **C** Flow cytometric analysis of CD56 in the shControl-, shFOSL2- and shEGR2-positive cells at days 28 and 35. **D** Quantification of CD56-positive cells in the GFP-positive cell population at days 28 and 35 (n=5). **E** Flow cytometric analysis of CD56 in the GFP-negative cell population at days 28 and 35. **F** Quantification of CD56-positve cells in the GFP-negative cell population at days 28 and 35 (n=5). **G** Quantification of CD11b-positive cells in the CD56^+^ cell population at day 35 (n=5). The results from five biological replicates are presented as the mean ± SEM. *P<0.05, **P<0.001, and ***P<0.0005.

In addition, we attempted to identify the signaling pathways through which FOSL2 and EGR2 were involved in driving NK cell differentiation by performing module map analysis of all the differential peaks. 18 modules were identified according to the patterns of their accessibility (**Table S3**), several of which were significantly enriched with known TFs that regulate NK cell differentiation (**Figure S6B**), including FOSL2 and EGR2, and many genes in the JAK-STAT pathway, such as the STAT family, IRF2, TBX21, and GATA3. These results suggest that FOSL2 and EGR2, may regulate NK cell differentiation through the JAK-STAT pathway (**Figure S6C**).

### Transcriptional regulatory network dynamics during NK cell differentiation

The dramatic chromatin accessibility differences during NK cell differentiation prompted us to check the time point-specific transcriptional regulatory network. Although some TFs have been discovered to regulate NK cell differentiation, little is known about the dynamics of the entire regulatory network during this process. Since the TF footprint pattern from the ATAC-seq reads can simultaneously directly reveal the binding profiles of hundreds of TFs with known cognate motifs [22], together with gene expression profiles, we can construct a regulatory network at each time point and assess how it changes during NK cell differentiation. First, we used HOMER to identify enriched TFs that bound to the cluster I and cluster II peaks shown in **Figure 3A** (P<0.05). We then examined the gene expression profiles of these TFs and found that 120 TFs were expressed in at least one stage during the differentiated process. By applying differential analysis, we obtained 14-26 TFs that were distinctly expressed at each stage (fold change>1.5) (**Figure S7**), and defined them as the nodes of the regulatory network. The connections (edges) between any 2 TFs were defined as follows: If the TF A motif is on the promoter of TF B, then we say TF A regulates TF B and draw an arrow from TF A to TF B. Here, only TFs that were expressed at the specific time point were consideration [58]. Using this method, we constructed the transcriptional regulatory network at each time points with both the enrichment (P-value) and expression information for all the relevant TFs (**Figure 5A-E**). Interestingly, day 7 specific TFs were densely interconnected at the beginning, and quickly vanished after two weeks (**Figure 5A**). In contrast, the day 35 specific network gradually grew out through the induction of relevant TFs. Many known regulators, such as EOMES, TBX21, ETS1, PRDM1, and GATA3 and also FOLS2 and EGR2 were increasingly enriched on the network (**Figure 5E**). Similar phenomena were also observed on other networks (**Figure 5B-D**). We believe the dynamics of the transcriptional regulatory network explain the increase in the proportion and the differentiation of NK cells.

**Figure 5.**
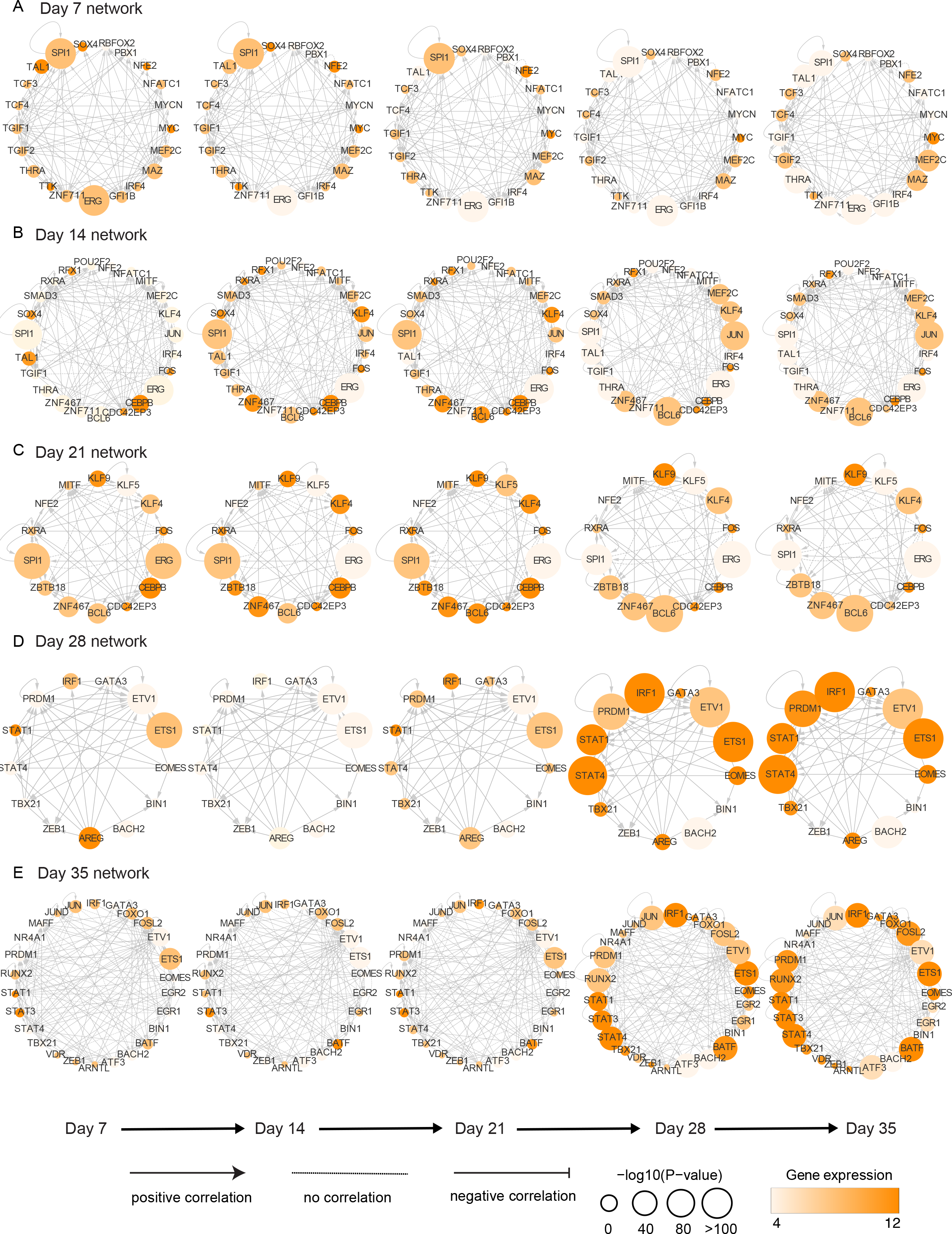
Transcriptional regulatory eetwork dynamics during NK cell differentiation. **A∼E** Cis-regulatory networks between TFs (nodes) enriched in at least one gene set and specifically expressed (fold change>1.5) at day 7 (**A**), day 14 (**B**), day 21 (**C**), day 28 (**D**) and day 35 (**E**). The arrow at the edge from node A to node B indicates that TF A regulates TF B by binding to the promoter site of the latter. The color of each node indicates the expression value of the gene encoding that TF, and the size of the circle represents the significance of the motif enrichment according to the P-value. The types of edges indicate the Pearson correlation between the gene expression profiles of the connected TFs: positively correlated (correlation coefficient>0.4); negatively correlated (correlation coefficient<-0.4); no correlation (correlation coefficient between −0.4 to 0.4).

## Discussion

NK cells are important innate immune cells that have been recognized as efficient effector cells to treat tumors. To better understand the differentiation machinery of NK cells and identify new regulators in the process, we studied the landscape of active elements during NK cell differentiation using the sensitive ATAC-seq method. As shown by the ATAC-seq results, three distinct clusters of DNA accessible elements were found. Further analysis showed that the chromatin accessibility correlated well with the level of the expression of the corresponding genes. In short, this study provides an epigenomic landscape and dynamics of NK cell differentiation and presents foundational profiles for studying the relationship between chromatin accessibility, gene expression and cell growth during this process (**Figure 2**).

TFs bind to their motifs and are often obligate nucleosome evictors and the creators of accessible DNA sites, and therefore we can use ATAC-seq to predict critical regulators in NK cell differentiation [26]. By motif analysis in HOMER, we found that several known TFs were enriched at different stages during NK cell differentiation. Similar results were also obtained using Genomica (**Figure 3B**). The discovery of known regulators strongly suggested the reliability of our analysis Furthermore, by integrating results from HOMER, Genomica and motif footprint analysis, we identified two novel TFs, FOSL2 and EGR2 that were essential for NK cell maturation. Knockdown of either of these two TFs significantly inhibited NK cell transformation in the *in vitro* NK cell differentiation system. Module map analysis suggested these two TFs may regulate NK cell through the JAK-STAT pathway, and therefore further studies of this pathway may facilitate the generation of NK cells and thus promote the NK cell-based immunotherapy. Overall, this study also provided a framework to identify new regulators from chromatin accessible data for NK cell differentiation.

TFs do not usually function alone, they always interact with other molecules to fulfill their unique roles. Hence, we depicted the transcriptional regulatory networks at different stages during NK cell differentiation. In order to construct a stable transcriptional regulatory network, we performed a rigorous screening of TFs to avoid stochastic fluctuations, and integrated both the enrichment (P-value) and expression information for all the relevant TFs. Therefore, even a change of either the enrichment or gene expression cutoff may result in different networks, the most critical TFs to the regulatory process still remain. However, since the screening of TFs mainly rely on the gene microarray, which is not as accurate as RNA-seq, the structure of these predicted network may not as robust as well.

From our previous study, we noticed that with a minimal cytokine cocktail, we can generate sufficient number of functional NK cells that express the cytokines necessary for NK cells and have a high effect on tumor cells [19]. Although there may also be a certain proportion of other lymphocytes, *in vitro* expansion of NK cells from peripheral blood (PB) or UCB cells has been successfully performed and developed in several clinical strategies to treat cancer [4, 5, 59-61]. Therefore, a comprehensive understanding of the regulatory patterns at each differentiated stage of the *in vitro*-derived NK cells in this system will help to uncover the underlying mechanisms of NK cell differentiation. The transcriptional regulatory network revealed in this study will lay the foundation for faster and better *in vitro* production of effective NK cells, thus facilitating NK cell-based immunotherapy. Moreover, we have identified two novel TFs, FOSL2 and EGR2, as essential regulators in controlling NK cell maturation and function. We also predicted the potential signaling pathways in which these two TFs may be involved and illustrated the dynamics of the transcriptional regulatory network during NK cell differentiation. In spite of the advantages of our strategy, there are two main limitations of this study: (1) Although induced NK cells are very similar to those produced *in vivo*, these two types of NK cells are still not identical. We observed certain differences between the induced NK cells and the NK cells produced *in vivo* in terms of chromatin accessibility (data not shown). (2) Before NK cells are fully developed, there is always a mixture of cell populations with stem cells, NK progenitor cells, immature NK cells and mature NK cells and other cell types that we are incapable to delineate at this moment, i.e. any bulk cell based analysis may neglect the huge heterogeneity between cells by default. Therefore, to fully uncover the regulatory mechanism, single cell technologies are required in the future to further delineate the cell-to-cell heterogeneity and regulatory dynamics at single cell precision.

## Materials and Methods

### Samples

Umbilical cord blood (UCB) samples were collected at birth from women with normal, full-term deliveries at Anhui Provincial Hospital, Hefei, after receiving informed consent. Approval was obtained from the Ethics Committee of the University of Science and Technology of China. The culture procedure for NK cell differentiation from UCB CD34^+^ cells has been previously described [19].

### Immunofluorescence staining

The cultured cells were post-fixed in 4% paraformaldehyde, blocked with 10% goat serum and stained with primary antibodies at 4°C overnight. Secondary antibodies were stained at 37°C for 1 h. Cell nuclei were stained with DAPI for 5 min at room temperature. Confocal images were acquired using a Zeiss LS710 microscope.

### RNA isolation and real-time PCR

Cultured cells were lysed in TRIzol reagent (Invitrogen), and total RNA was extracted following the manufacturer’s instructions. cDNA was synthesized with Moloney murine leukemia virus reverse transcriptase (Invitrogen) and oligo (dT) 20 primers. Then, SYBR Premix Ex Taq (TaKaRa) was used for real-time PCR (primer sequences, **Table S4**). The data were analyzed using the 2-ΔΔCt method.

### Lentivirus production and transduction

To produce lentiviral particles, 293T cells were transfected with the plasmids PLKO.1, pRRE, pREV and pVSV-G via Lipofectamine 2000 (Invitrogen) following the manufacturer’s protocol. Then, we harvested the supernatants 48 and 72 h post-transfection. To remove cell debris, supernatants were centrifuged (3000 rpm, 10 min), and then, the lentivirus particles were concentrated by ultracentrifugation at 50000 g for 2 h at 4°C. Finally, the virus particles were gently resuspended in HBSS and stored at −80°C. After UCB CD34^+^ cells were cultured with multiple cytokines for 14-18 d, we incubated the lentivirus and the cultured cells with polybrene (5 µg/ml) and centrifuged them at 1000 rpm for 70 min at 10°C.

### Statistical analysis

Statistical significance was analyzed using unpaired two-tailed t-tests. P-values less than 0.05 were considered statistically significant.

### Microarray

The microarray was performed using an Affymetrix GeneChip® PrimeView Human Gene Expression Array. The signal values of the samples were normalized using RMA. Differential analysis was performed using Student’s t-test as previously described [19]. The microarray data were under accession number GEO: GSE87787.

### ATAC-seq

ATAC-seq was performed as previously described [22], and 2×150 paired-end sequencing was performed on an Illumina HiSeq X-10 to yield, on average, 78 M reads/sample.

### ATAC-seq analysis

Sample reads from biological replicates were then grouped together and divided into 8 categories: day 7, day 14, day19, day 21, day 24, day 26, day 28 and day 35. Intrinsic analysis and other ATAC-seq analysis was performed same as our previous work [26, 62].

### Difference analysis

All samples were grouped into 8 categories (16 samples): day 7, 14, 19, 21, 24, 26, 28 and day 35. Data normalization and significance analysis were performed via pairwise comparison between the 8 categories using DESeq2 [63], with a P<0.01, log2-fold change>5, FDR<0.01, and intrinsic analysis [26] and with a z-score>1. We finally obtained 6401 differential peaks. Unsupervised clustering was performed using Cluster 3.0 and visualized in Treeview. GREAT [33] was used to predict enriched functions and Gene Ontology.

### Stage-specific peaks

Each stage (e.g., day 7) consists of genes that were induced (>1.5-fold change) at that stage compared with all the other stages. We defined peaks that were accessible only in one stage as stage-specific peaks, and those that were accessible at all stages as conservative peaks.

### Define promoter

We define the range of 2kb around the transcription start site as a promoter.

### TF motif enrichment analysis

HOMER [29] was used to perform the TF enrichment analysis with the following options: findmotifs.pl input.fa fasta output/ for **Figure 3A** and findmotifs.pl input.fa human uotputdir -fasta bg.fa for **Figure S5A**. TF enrichment analysis using Genomica and TF footprinting analysis was performed the same as our previous work [26].

### Gene Module Analysis

Gene module analyses were performed using WGCNA [64] in **Figure S6B** with the options SoftPower = 20, minModuleSize = 30.

### STRING analysis

Protein-protein interaction analyses were performed using STRING [65] in **Figure S6C**.

### Transcriptional regulatory network

We used HOMER to find the transcription factors that bind to cluster I and cluster II and obtained a total of 253 transcription factors that could regulate these differentially expressed elements (P<0.05), of which 120 were expressed in the microarray profile of the corresponding sample. If TF A bound to the promoter of TF B, we defined TF A as a regulator of TF B and then constructed a transcriptional regulatory network. Networks of TFs were assembled from this set of 120 TFs that were expressed in at least one sample. An edge between TF A and TF B indicated that TF A bound to the promoter of TF B [58].

## Data Availability

### GSA accession

The raw sequence data have been deposited in the Genome Sequence Archive [66] in BIG Data Center [67], under the accession number CRA000846 and are publicly accessible at http://bigd.big.ac.cn/gsa/s/u7fdeNV3.

## Acknowledgements

This work was supported by the National Natural Science Foundation of China grant 81788101 (to Z.T.), the National Key R&D Program of China 2017YFA0102903 (to K.Q.), the National Natural Science Foundation of China grant 91640113 (to K.Q.) and 31771428 (to K.Q.), the key project of Natural Science Foundation of China 81330071 (to H.W.). We thank the USTC supercomputing center and the School of Life Science Bioinformatics Center for providing supercomputing resources for this project.

## Authors’ contributions

KQ and HW conceived the project, YW and YL performed all the cell sorting and ATAC-seq library experiments, KL performed most of the data analysis, QY analyzed the microarray data, HW and ZT discussed the results and provided advices, KL and KQ wrote the manuscript.

---

**Supplementary Figure S1. Landscape of DNA accessibility during NK cell differentiation**

**A** Fractions of CD56^+^CD3^−^ cells in the total gated cells during a 35-day time course.

**B-C** Quality control analysis of ATAC-seq data. **B**: The TSS enrichment score for all samples. **C**: The fragment length distribution of all the mapped reads.

**D** Known cell surface markers: Normalized ATAC-seq profiles of the *CD34*, *KIT*(CD117), *KLRD1*(CD94) and *NCAM1*(CD56) gene loci at different stages during NK cell differentiation.

**E** Normalized ATAC-seq profiles of the *EOMES* (left) and *GATA3* (right) gene loci at different stages during NK cell differentiation. These genes are known to regulate NK cell differentiation, and ATAC-seq signals were obtained from the UCSC genome browser.

**Supplementary Figure S2.** T**he correlation analysis on the samples**

A The Pearson correlation analysis on the replicates at each time point. R at the top is the Pearson correlation coefficient.

B Heatmap of the Pearson correlation between all the samples with unsupervised clustering performed in Cluster 3.0.

**Supplementary Figure S3. Epigenomic signatures of NK cell differentiation at different stages**

**A** Venn diagram of peaks identified at each stage of NK cell differentiation. Specific peaks were defined as peaks that were identified only at a specific time point, and conserved peaks were those identified at all stages during the process.

**B** The top most significant GO terms of all the stage-specific and conservative peaks.

**Supplementary Figure S4. Epigenomic signatures of different stages during NK cell differentiation**

**A** Heatmap of all the 6401 differential regulatory elements in all the samples. Each column is a sample, and each row is a peak. Samples and peaks were organized by two-dimensional unsupervised hierarchical clustering. The color scale shows the relative ATAC-seq signal intensities as indicated. Top: samples at all time points were categorized into three groups, early stage: days 7∼21 (orange); interim stage: days 24∼28 (yellow) and late stage: days 28∼35 (green). Samples from the same cluster are labeled with the same color. Left: differential peaks are categorized into 3 clusters.

**B** Distance from all the peaks in cluster I, II, and III to their nearest genes. Known TFs regulating NK cell differentiation are labeled.

**C** Principle component analysis of chromatin accessibility during NK cell differentiation. Three clusters were also identified: early (days 7∼21), interim (days 24 ∼28), and late (day 35).

**Supplementary Figure S5**. **Transcription factor occupancy network during NK cell differentiation**

**A** The top ten TF motifs enriched in cluster I (left) and cluster II (right) peaks, with enrichment P-values estimated from HOMER. TFs known to regulate NK cell differentiation are color-coded.

**B** Left: heatmaps of the gene expression profiles of the genes predicted to be regulated by RUNX, T-bet, EGR2 and FOSL2. Unsupervised hierarchical clustering was performed. Right: the top ten most significant GO terms enriched in up-regulated (orange) and down-regulated (green) genes predicted to be regulated by each TF on the left.

**Supplementary Figure S6. EGR2 and FOSL2 are necessary for NK cell differentiation**

**A** Real-time qPCR analysis (n=3) of the genes *EGR2* (left) and *FOSL2* (right). The results from three replicates are presented as the mean ± SEM.

**B** Module map analysis in Genomica: representative modules from module map analysis in Genomica. Top: the most significantly enriched TFs in each module; bottom: the chromatin accessibility changes of the peaks in the corresponding module compared with day 7. Right: genes associated with peaks in that module.

**C** Signaling pathways of known and predicted TFs that regulate NK cell differentiation.

**Supplementary Figure S7. Define time-specific TFs based on differential expression analysis**

A: Distribution of all TF’s fold change. X axis represents the fold change. y axis represents the density of TF under different fold change. Std indicates Standard Deviation. Foldchange1.5 is 2 standard deviations away. Colors represent different time points.

B: The number of TFs that qualify fold changes. X axis represents different FDC (fold change). The y axis represents the number of TFs that pass the fold change cutoff.

C: The ratio of known TFs versus all known TFs that qualify the different fold change cutoffs. Known TFs: TFs regulating NK cell development reported in the literature.

