## Supplemental figures for "Landscape and dynamics of the transcriptional regulatory network during natural killer cell differentiation"

Figure S1

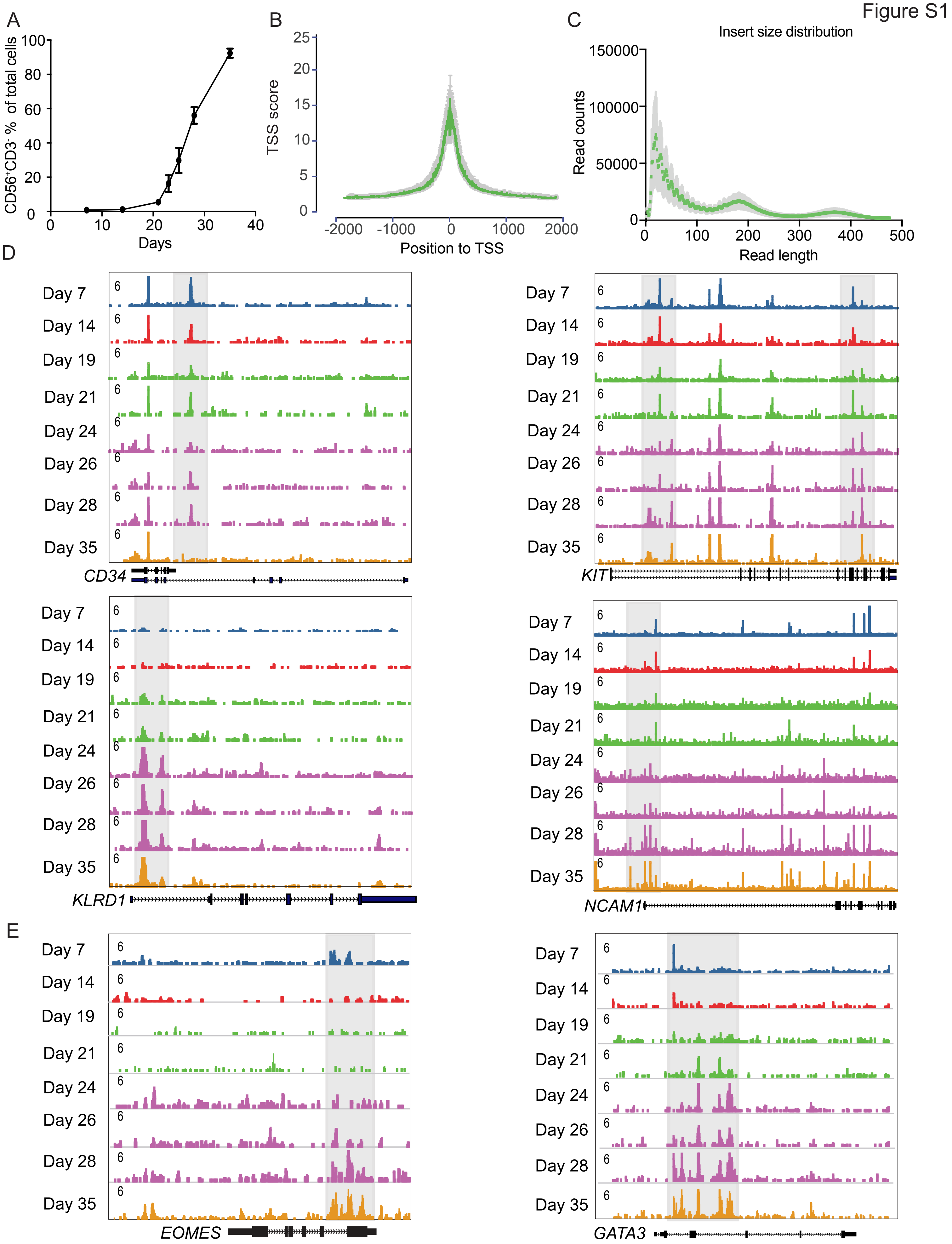


**Supplementary Figure S1. Landscape of DNA accessibility during NK cell differentiation**

A: Fractions of CD56$^+$CD3$^-$ cells in the total gated cells during a 35-day time course.

B-C: Quality control analysis of ATAC-seq data. B: The TSS enrichment score for all samples.

C: The fragment length distribution of all the mapped reads.

D: Known cell surface markers: Normalized ATAC-seq profiles of the CD34,KIT(CD117),KLRD1(CD94) and NCAM1(CD56) gene loci at different stages during NK cell differentiation.

E: Known TFs: Normalized ATAC-seq profiles of the EOMES (left) and GATA3 (right) gene loci at different stages during NK cell differentiation. These genes are known to regulate NK cell differentiation, and ATAC-seq signals were obtained from the UCSC genome browser.

Figure S2

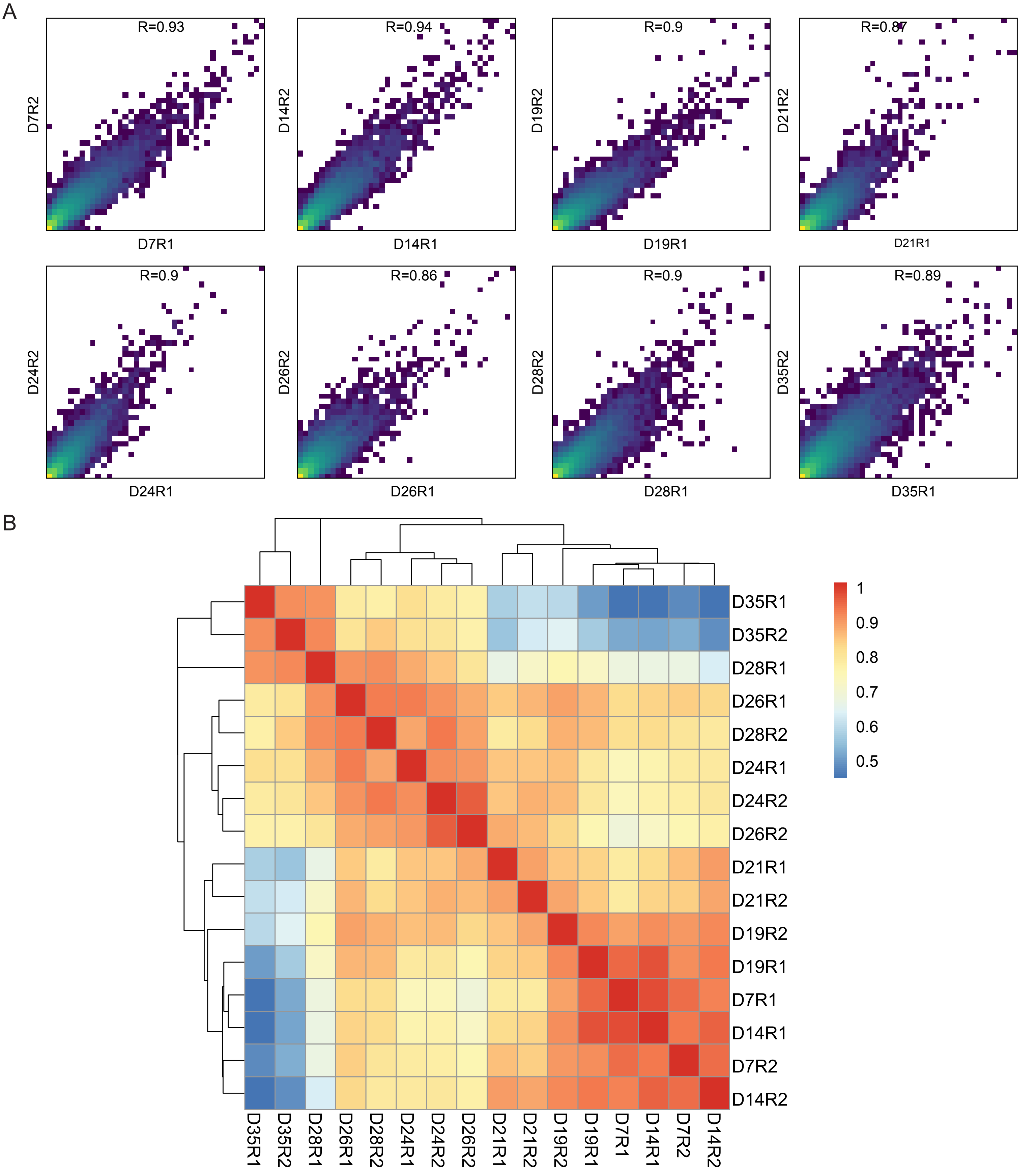


**Supplementary Figure S2. The correlation analysis on the samples**

A: The correlation analysis on the replicates at each time point. R at the top is the pearson correlation.

B: Heatmap of the Pearson correlation between all the samples with unsupervised clustering performed in Cluster 3.0.

Figure S3

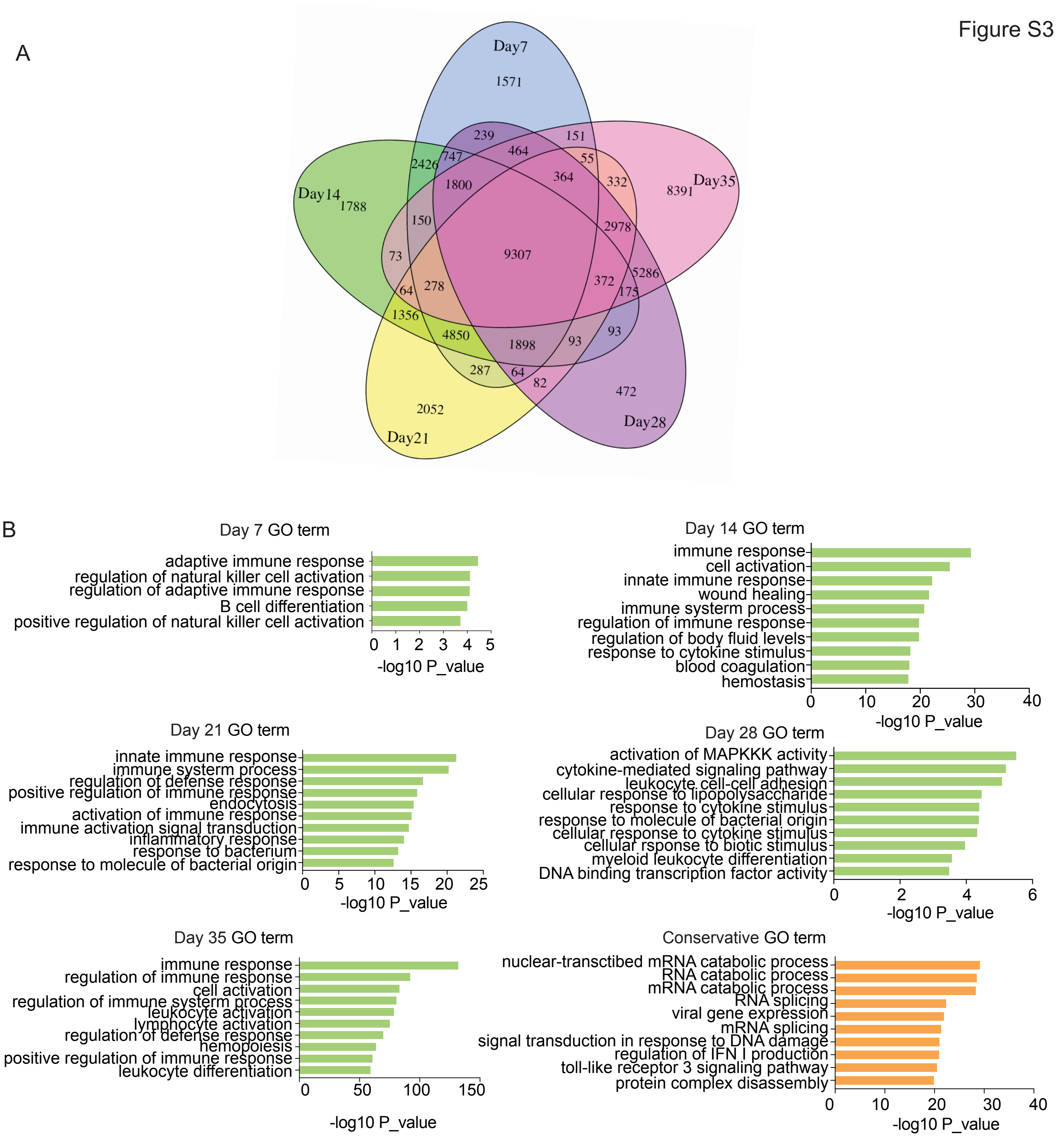


**Supplementary Figure S3. Epigenomic signatures of NK cell differentiation at different stages**

A: Venn diagram of peaks identified at each stage of NK cell differentiation. Specific peaks were defined as peaks that were identified only at a specific time point, and conserved peaks were those identified at all stages during the process.

B: The top most significant GO terms of all the stage-specific and conservative peaks.

Figure S4

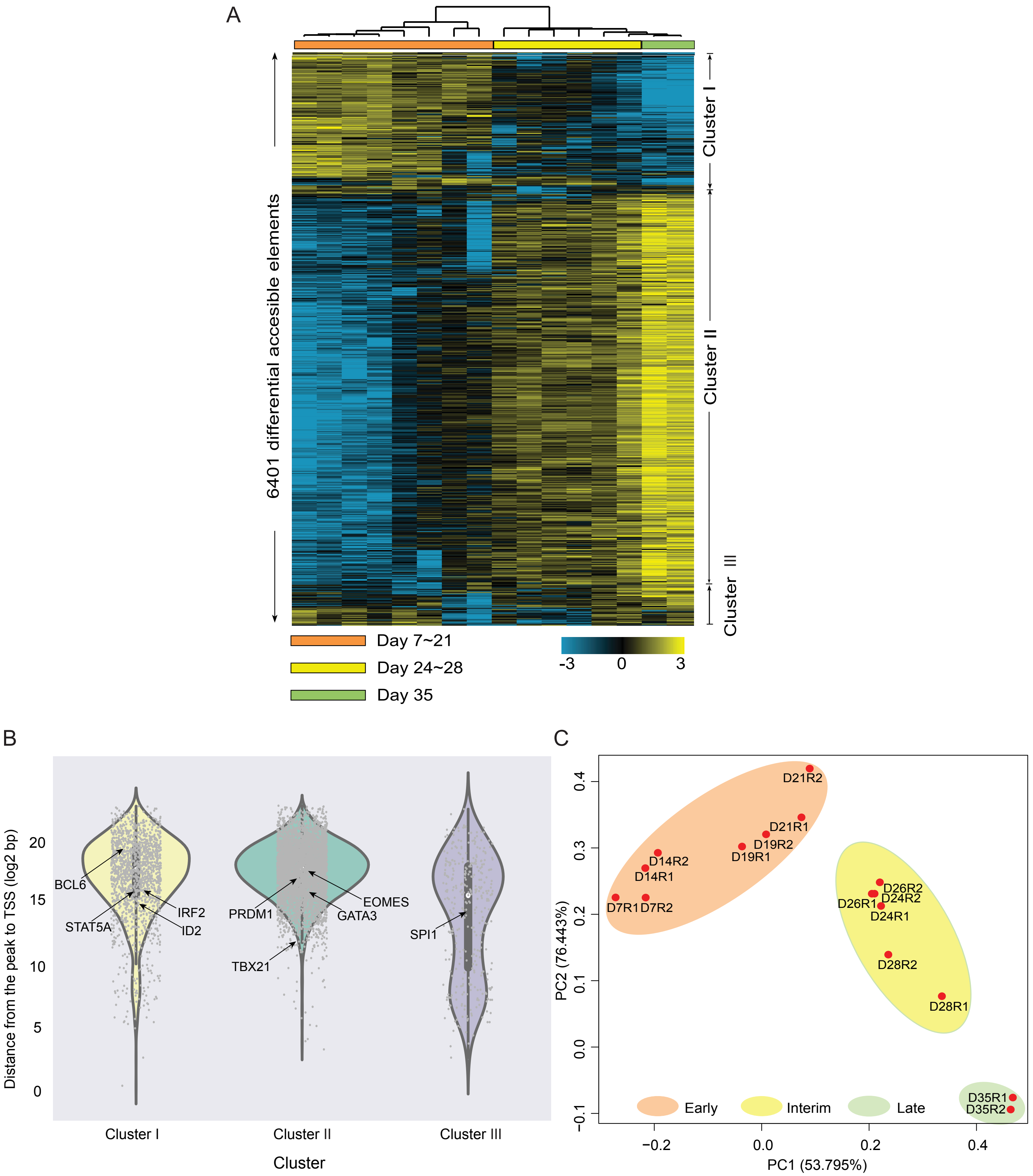


**Supplementary Figure S4. Epigenomic signatures of different stages during NK cell differentiation**

A: Heatmap of all the 6401 differential regulatory elements in all the samples. Each column is a sample, and each row is a peak. Samples and peaks were organized by two-dimensional unsupervised hierarchical clustering. The color scale shows the relative ATAC-seq signal intensities as indicated. Top: samples at all time points were categorized into three groups, early stage: days 7~21 (orange); interim stage: days 24~28 (yellow) and late stage: days 28~35 (green). Samples from the same cluster are labeled with the same color. Left: differential peaks are categorized into 3 clusters.

B: Distance from all the peaks in cluster I, II, and III to their nearest genes. Known TFs regulating NK cell differentiation are labeled.

C: Principle component analysis of chromatin accessibility during NK cell differentiation. Three clusters were also identified: early (days 7~21), interim (days 24 ~28), and late (day 35).

Figure S5

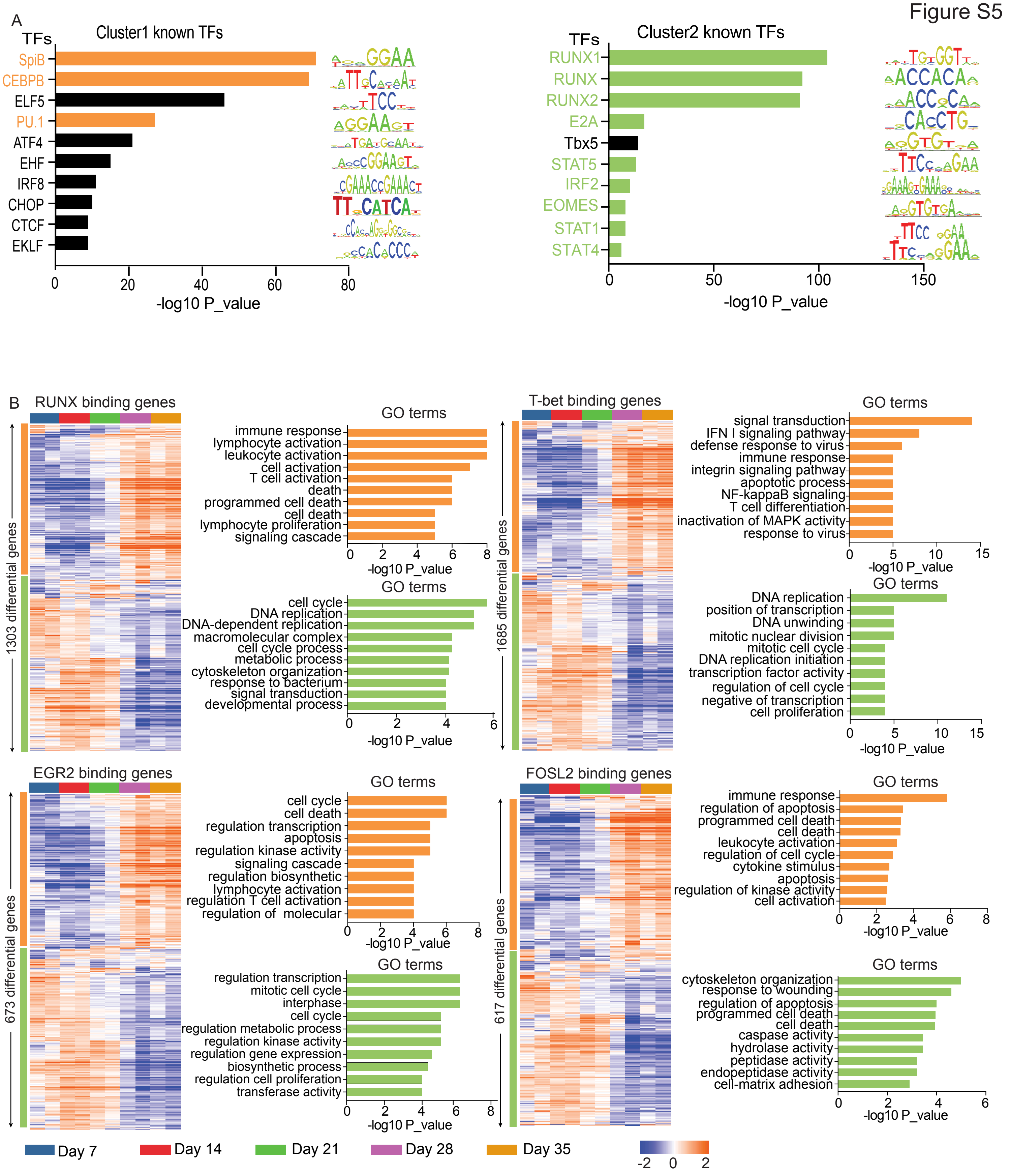


**Supplementary Figure S5.Transcription factor occupancy network during NK cell differentiation**

A: The top ten TF motifs enriched in cluster I (left) and cluster II (right) peaks, with enrichment P-values estimated from HOMER. TFs known to regulate NK cell differentiation are color-coded.

B: Left: heatmaps of the gene expression profiles of the genes predicted to be regulated by RUNX, *T-bet*, EGR2 and FOSL2. Unsupervised hierarchical clustering was performed. Right: the top ten most significant GO terms enriched in up-regulated (orange) and down-regulated (green) genes predicted to be regulated by each TF on the left.

Figure S6

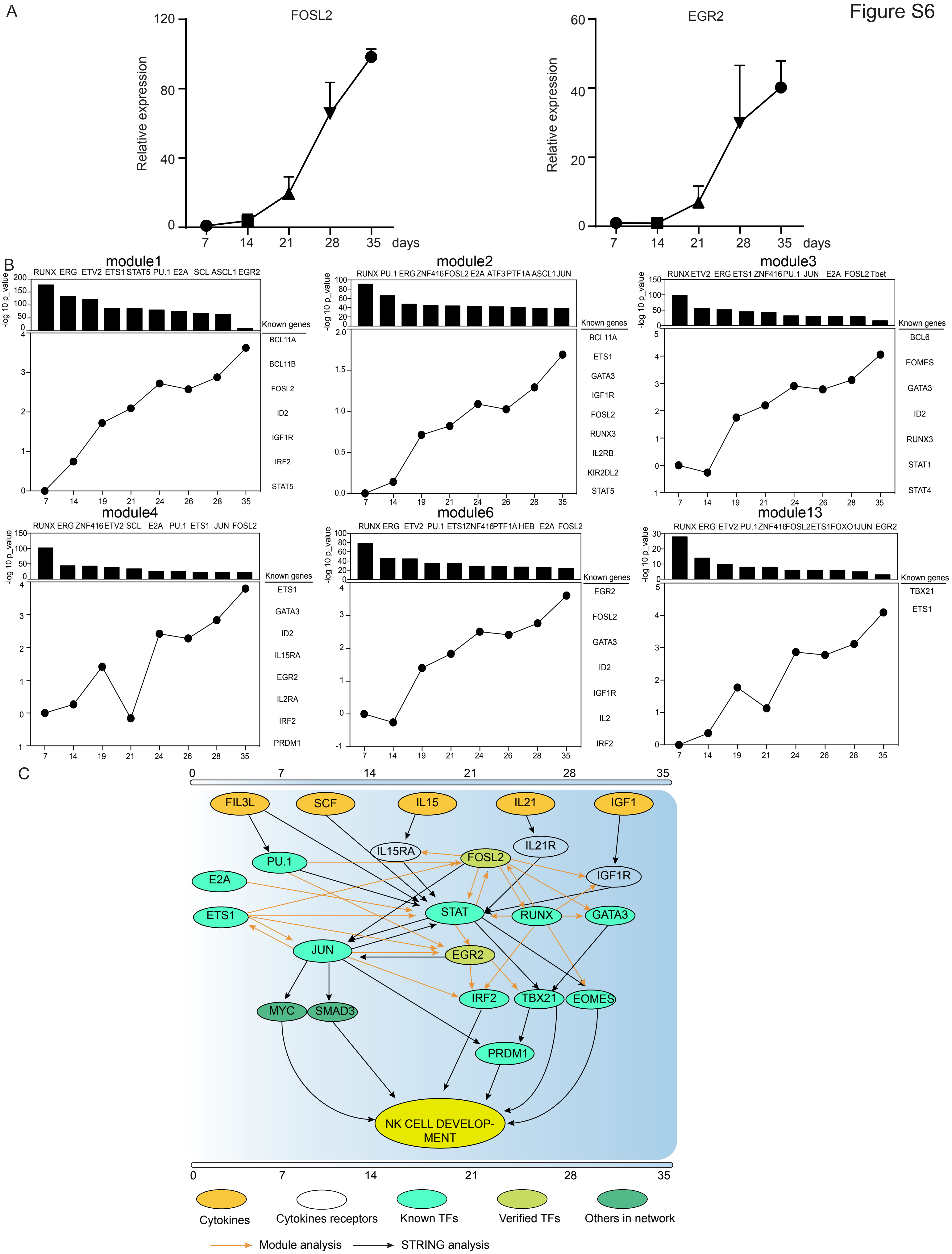


**Supplementary Figure S6. FOSL2 and EGR2 are necessary for NK cell differentiation**

A: Real-time qPCR analysis (n=3) of the genes EGR2 (left) and FOSL2 (right). The results from three replicates are presented as the mean ± SEM.

B: Module map analysis in Genomica: representative modules from module map analysis in Genomica. Top: the most significantly enriched TFs in each module; bottom: the chromatin accessibility changes of the peaks in the corresponding module compared with day 7. Right: genes associated with peaks in that module.

C: Signaling pathways of known and predicted TFs that regulate NK cell differentiation.

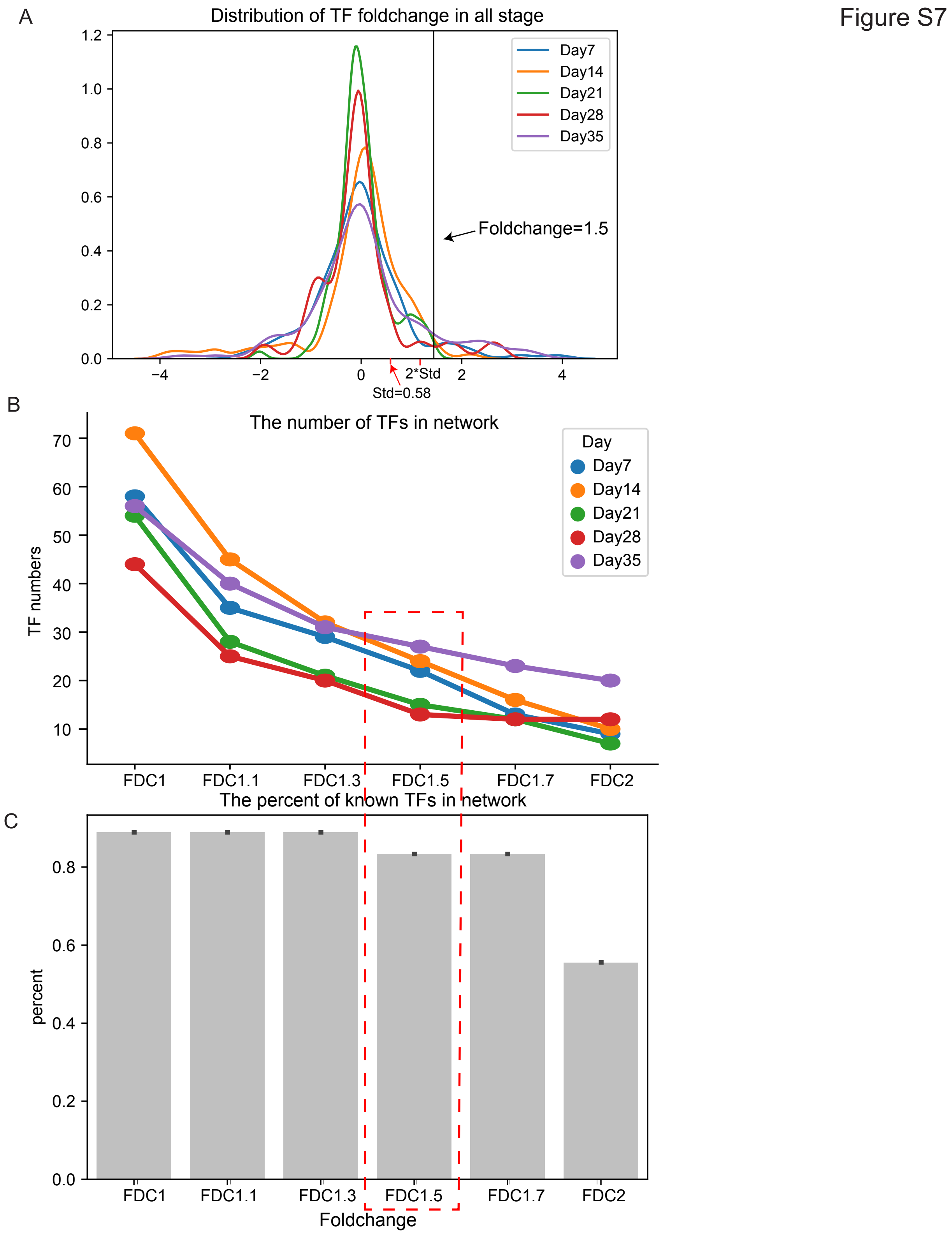


**Supplementary Figure S7.Define period-specific TFs based on differential expression of TFs**

A: Distribution of all TF's foldchange. X axis represents different foldchange, y axis represents the density of TF under different foldchagne. Std indicates Standard Deviation. Foldchange1.5 is outside the std of twice.Colors represent different times。

B: The number of TFs that satisfy different foldchanges. X axis represents different FDC(foldchange). The y axis represents the number of TFs

C: The ratio of known TFs satisfying different foldchanges to all known TFs.Known TFs :TFs regulating NK cell development reported in the literature.
